# *transmorph*: a unifying computational framework for single-cell data integration

**DOI:** 10.1101/2022.11.02.514912

**Authors:** Aziz Fouché, Loïc Chadoutaud, Olivier Delattre, Andrei Zinovyev

## Abstract

Data integration of single-cell data describes the task of embedding datasets obtained from different sources into a common space, so that cells with similar cell type or state end up close from one another in this representation independently from their dataset of origin. Data integration is a crucial early step in most data analysis pipelines involving multiple batches and allows informative data visualization, batch effect reduction, high resolution clustering, accurate label transfer and cell type inference. Many tools have been proposed over the last decade to tackle data integration, and some of them are routinely used today within data analysis workflows. Despite constant endeavors to conduct exhaustive benchmarking studies, a recent surge in the number of these methods has made it difficult to choose one objectively for a given use case. Furthermore, these tools are generally provided as rigid pieces of software allowing little to no user agency on their internal parameters and algorithms, which makes it hard to adapt them to a variety of use cases. In an attempt to address both of these issues at once we introduce *transmorph*, an ambitious unifying framework for data integration. It allows building complex data integration pipelines by combining existing and original algorithmic modules, and is supported by a rich software ecosystem to easily benchmark modules, analyze and report results. We demonstrate *transmorph* capabilities and the value of its expressiveness by solving a variety of practical single-cell applications including supervised and unsupervised joint datasets embedding, RNA-seq integration in gene space and label transfer of cell cycle phase within cell cycle genes space. We provide *transmorph* as a free, open source and computationally efficient python library, with a particular effort to make it compatible with the other state-of-the-art tools and workflows.

## Introduction

Batch effects occur in most applications involving datasets gathered across multiple sources or experiments, and describe strong dataset-specific signals which are often irrelevant to the studied questions. Data integration is a computational paradigm aiming to learn a joint embedding of datasets in which batch effects are regressed out [Fig. 1a], so that only dataset-independent factors are expressed. The idea is to learn this space by leveraging information contained in several datasets, each of those being biased by its own specific batch effects. We focus here on the so-called *horizontal data integration* [1] which seeks to integrate cells measured in the same domain with overlapping feature spaces. This is different from *vertical* and *diagonal data integration* where cells are measured in different domains, also known as multi-omics data integration. This scenario involves specific strategies and algorithms which are beyond the scope of this work (see for instance [2]).

**Figure 1:**
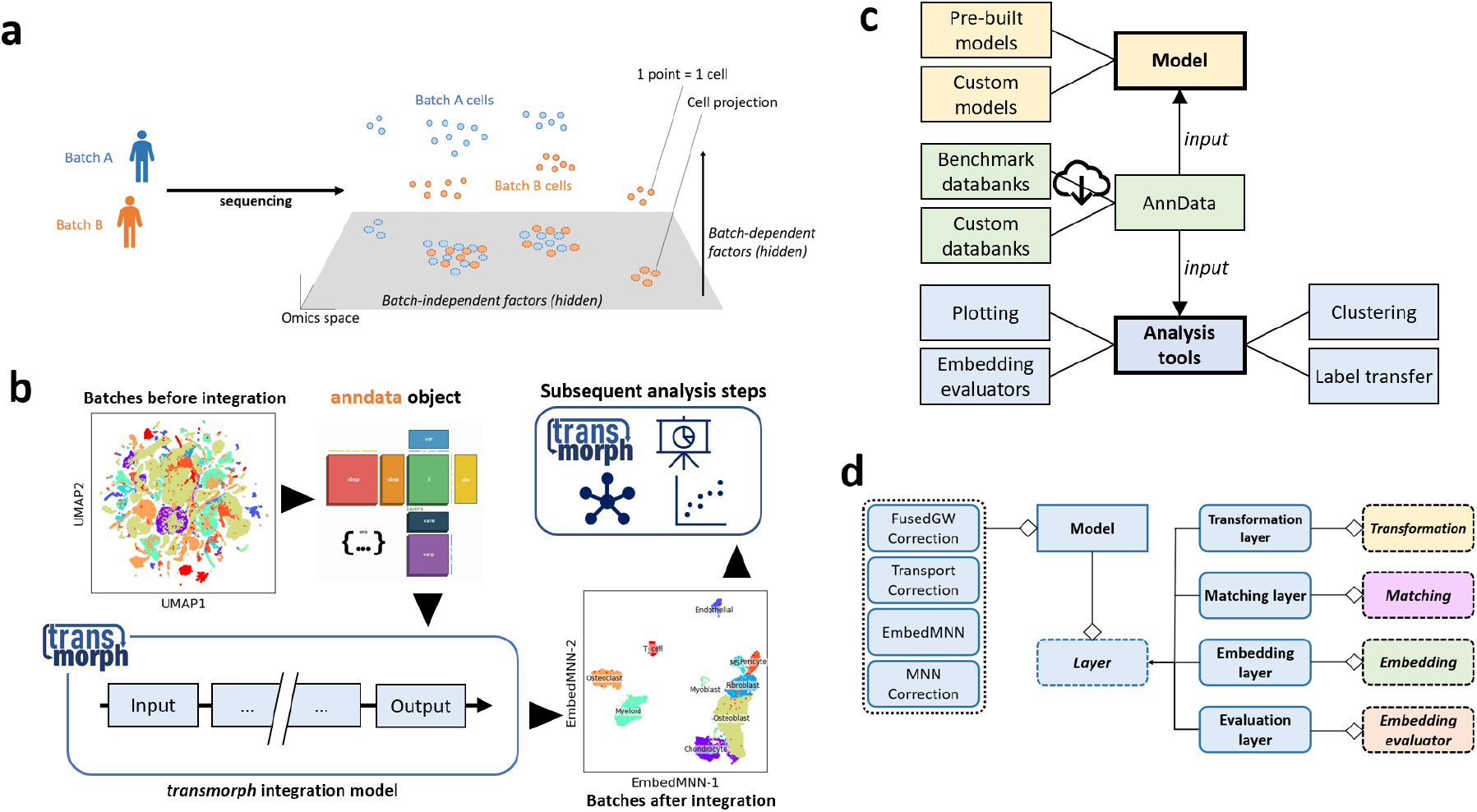
**a.** Schematic representation of the data integration problem, consisting in learning a batch-independent embedding of the datasets. **b.** Usage of *transmorph* models and software environment in a python single-cell analysis pipeline. **c.** *transmorph* ecosystem, with model API and analysis tools. **d.** *transmorph* model API architecture used to build complex data integration models.

Data integration is an important preprocessing step for applications involving several datasets. In some cases something as simple as centering and normalization/scaling of features may suffice, but more complex batch effects often require more subtle, dedicated algorithms to be satisfyingly removed. Data integration can serve various purposes. The most common usage is to embed items from all datasets into a joint low dimensional space like in the Harmony tool [3], which can then be used to carry out downstream learning algorithms such as clustering, label transfer or visualization. Another use case is to directly perform integration in the initial domain like in the MNN tool [4], so that algorithms needing interpretable features such as matrix factorization methods can be used. Finally, integration can be carried out without embedding data points into an explicit feature space, for instance by outputting a joint graph of cells across datasets like in the BBKNN tool [5].

Data integration finds particularly important applications in single-cell biology. A single-cell dataset is generated from a biological tissue, and contains individual molecular measurements (for instance gene expression, SNPs or chromatin accessibility) about single cells of the tissue. The strength of single-cell analysis is its ability to both provide an insight into intrinsic cell state, while also giving access to population-level information, that can for instance be used to estimate cell types distribution within a tissue. Due to genetic and environmental differences between individuals, batch effects are very prone to appear in medicine when dealing with single-cell datasets coming from different patients. The intertwine of batch-dependent and batch-independent factors is an obstacle for the analysis of large comprehensive datasets built by aggregating data from different individuals, notably when building cell atlases (see for instance [6]). Data integration is therefore a necessary technology to develop in order to mitigate dataset-specific signal while preserving relevant biological signal proper to the system of interest.

Many approaches have been proposed over the last decade to tackle data integration in single-cell, resorting to a variety of algorithmic strategies. Some methods such as mutual nearest neighbors (MNN) [4], CONOS [7], Harmony [3], Seurat [8] and BBKNN [5] work under the hypothesis that batch effects are almost orthogonal to biological effects. This allows these tools to leverage data geometry using nearest neighbors across datasets as a proxy for cell-cell similarity. Other data integration methods like SCOT [9] and Pamona [10] use optimal transport-based algorithms to estimate cell-cell similarity, assuming the existence of similar manifolds supporting datasets containing cells of the same type. We would also like to mention the existence of neural network-based tools like the ones implemented in scVI [11], which make use of variational autoencoder compression to directly learn dataset embeddings inside a joint latent space.

If the ever-growing number of data integration methods is beneficial by bringing new ideas to the field, and despite constant efforts of many talented researchers to provide exhaustive benchmarks like in [12, 13], it becomes more and more difficult to choose the right data integration methodology for a given application [1]. Furthermore, existing data integration tools are provided as rigid algorithmic pipelines, allowing little to no user agency to adapt them to specific use cases, for instance when the output type of a certain tool is not adapted to the needs of a given workflow. This urges the need for a data integration framework which would help conceiving data integration pipelines by articulating computational modules, factoring boilerplate code and common preprocessing steps, while providing a sound interface to work with existing workflows [Fig 1b] as well as benchmarking algorithms and datasets. To address these needs we introduce *transmorph* [Fig 1c], an open-source data integration framework implemented in python providing great modularity, high integration quality and performance, robust implementation, and the ability to be extended over time with new cutting-edge algorithms. From a computational point of view, *transmorph* works directly with AnnData [14] objects as input, which allows it to be natively used within any *scanpy* [15] workflow.

## Results

### *transmorph* allows conceiving end-to-end data integration models

Despite achieving good integration results in specific use cases, we believe that existing data integration algorithms are flawed by their intrinsic rigidity. By constraining the user to a fixed algorithm, they tend to excel in some use cases while struggling in others, as we show in the next sections. Subsequently, the lack of access to their internal algorithms makes results difficult to interpret. Also, these internal algorithms cannot be swapped for something more appropriate when needed, for instance when their output type does not fit the next algorithms’ input type.

To address these limitations we present *transmorph*, a novel ambitious data integration framework. It features a modular way to create data integration algorithms using basic algorithmic and structural blocks, as well as analysis tools including embedding quality assessment and plotting functions. The framework also provides annotated, high quality and ready-to-use datasets to benchmark algorithms [Fig 1c]. Finally, it is meant to be easily expansible by allowing the user to define new algorithmic modules if necessary. In this framework, data integration models can be assembled by combining four classes of algorithms: transformations, matchings, embeddings and checkings [Fig 1d].

- **Transformation** algorithms take as input a set of datasets and return a new representation for each of them, embedded in some feature space (there can be one separate feature space per dataset, or one common feature space). Transformations are generally used during preprocessing: classic examples are PCA, neighborhood-based data pooling or common highly variable genes selection.
- **Matching** algorithms estimate a similarity measure between cells across datasets. They are the core component of our integration framework, as their quality directly influences cell-cell proximity in the final embedding. *transmorph* uses three main categories of matching; (a) label-based matchings which require datasets to be labeled beforehand and matches items of similar label; (b) neighbor-based matchings which match items close items with respect to some metric; (c) transport-based matchings which leverage a distance metric between items within or across datasets to compute a similarity between items relying on topological correspondence. The next sections demonstrate how important it is to choose a matching which is relevant for a given application.
- **Embedding** algorithms are a special class of transformations which take as additional input similarity relationships between samples that were estimated via a matching. They return an integrated view of all datasets jointly embedded in a common feature space, so that matched items tend to be close from one another in the final representation. The embedding step is in general the last step in an integration model, and is chosen depending on the required output type. For instance, a joint embedding of datasets in an abstract space is suited for applications like visualization or clustering, while matrix factorization algorithms often require the embedding to be performed in an expressive feature space. The next sections show how choosing the right embedding algorithm allows adapting an integration pipeline to a given scenario.
- **Checking** algorithms are special quality control points that can be added to a pipeline in order to test a condition. They are used to either set a branching point that leads to different outcomes, or create an iterative structure within a model (“ *repeat until the integrated representation satisfies this property*”). This type of strategy is notably used within the Harmony algorithm, where an iterative clustering and correction procedure is applied until an integration metric (Local Inverse Simpson’s Index in this case) is considered to be satisfactory.

This expressive framework allows building complex data integration models suited for many applications, with high computational efficiency and integration quality because each algorithmic module can be optimized independently. It also provides an objective comparison between algorithmic modules for a given application. Finally, it is supported by a sound software ecosystem with benchmarking databanks, pre-built models and post-analysis tools which allows one to easily setup entire data integration pipelines [Fig 1c]. Our framework is provided as an open-source python package, and the next results showcase its capabilities to solve a variety of challenging real-life problems in single-cell, while being on par with existing tools in terms of performance. It has been developed to be easily used in notebook environments, with a strong focus on computational efficiency so that models can be run on small machines in a reasonable time, even in applications involving tens of thousands of cells and more than ten different datasets and cell types.

### Building a high quality model to tackle joint dataset embedding

The first data integration challenge consists in computing a low dimensional joint embedding of two or more datasets, so that similar cells end up close to one another independently from their source. This is typically used for visual data exploration or as a preprocessing step before carrying out a clustering algorithm, allowing clusters to only depend on cell type rather than on the original batch. A good joint dataset embedding algorithm should be able to function in a fully unsupervised fashion while being improved by additional labeling information, and should not require to choose a reference dataset as this induces an important bias. Ideally, it should also be able to tackle joint embedding of more than two datasets at the same time, with reasonable computational efficiency.

We selected three state-of-the-art algorithms able to solve this problem: Harmony [3] which uses a clustering-driven, iterative strategy to optimize the embedded representation. ScVI [11], a deep learning framework which uses variational autoencoders to compute a latent integrated representation of datasets. BBKNN [5] which builds a weighted joint graph of datasets together using a batch-balanced variant of *k*-nearest neighbors. We embedded Harmony, scVI latent representation and BBKNN result into a 2D space using UMAP [16] so that the output space is comparable between all methods. We also included the *transmorph* pre-built model EmbedMNN to the benchmark, described in [Fig 2b]. This integration model is inspired from CONOS [7], and combines a nearest neighbors-based joint graph construction step with a low dimensional graph construction, followed by an embedding step using UMAP or minimum distortion embedding (MDE) [17]. EmbedMNN can work either in a fully unsupervised fashion, or can take into account label information to prune matching edges between samples of different labels; we test both variants in this application.

**Figure 2:**
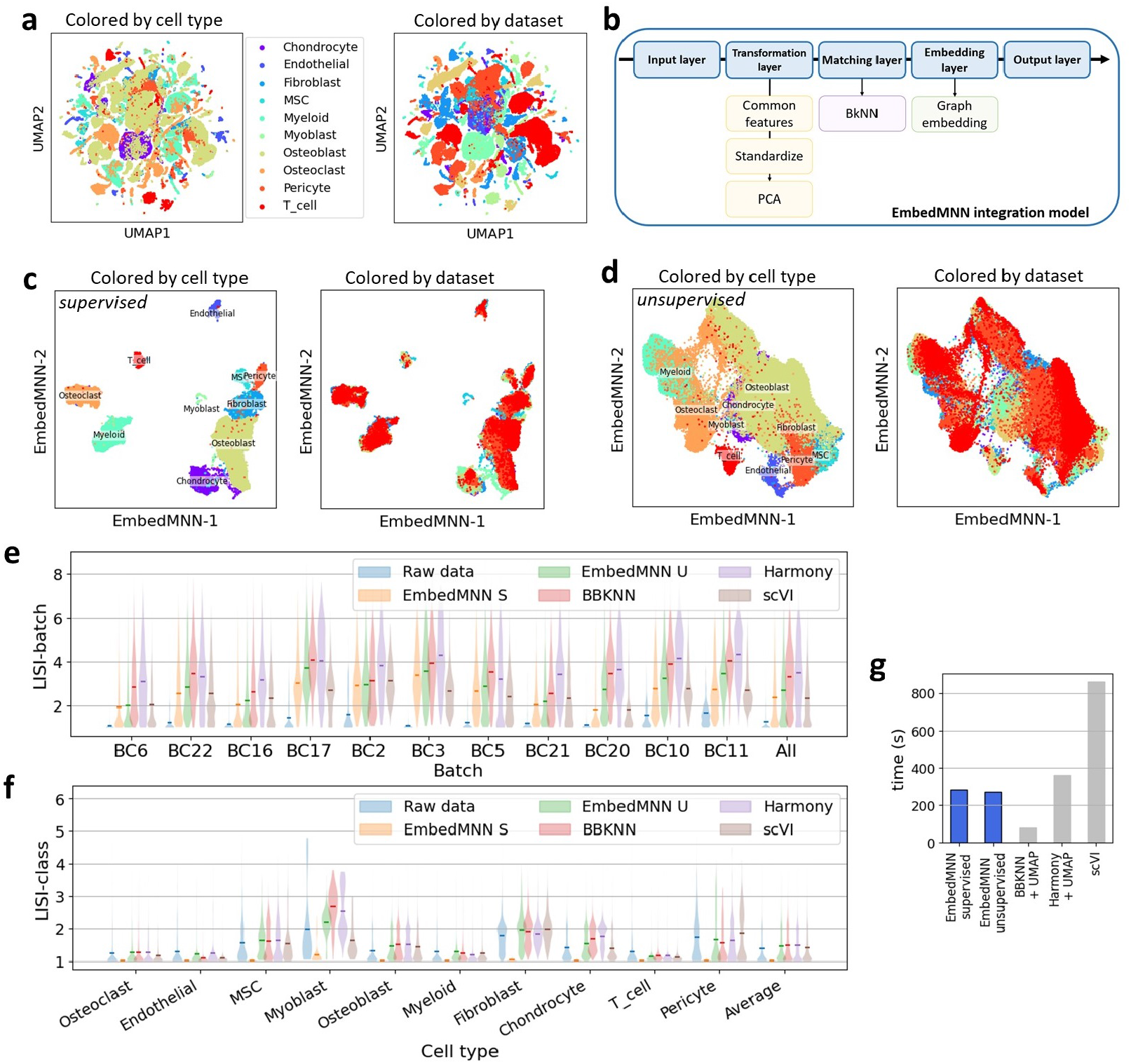
**a.** UMAP representation of the osteosarcoma datasets (n = 64, 557) in their common genes space. **b.** Architecture of the EmbedMNN integration model, computational modules are executed left to right and top to bottom. **c.** Integration results with the supervised version of EmbedMNN. **d.** Integration results with the unsupervised version of EmbedMNN. **e.** LISI-batch score of various integration algorithms (higher is better), mean is marked. **f.** LISI-class score of various integration algorithms (lower is better), mean is marked. **g.** Execution time of various integration algorithms.

The benchmarking databank consists of 11 single-cell osteosarcoma datasets gathered from [18], containing approximately 65,000 cells in total, which have been automatically annotated using 10 different cell types. This use case is quite challenging due to dataset size, number of batches and number of classes, but illustrates a reasonable real-life use case of data integration. Integration performance can be objectively measured through four integration metrics: batch and class mixing using a lightened version of local inverse Simpson’s index (LISI) introduced in Harmony, clustering specificity using Louvain or Leiden community detection algorithm [19, 20], and computation time. It is to note that Harmony directly uses batch-LISI as a criterion during its optimization procedure, so we have to expect it to have superior batch-LISI scores.

All methods were able to compute the integrated embedding in a reasonable amount of time given the number of data points [Fig 2g, Sup. Fig. 1], with the best performer being BBKNN + UMAP with 1min10s, taking advantage of the excellent C++ nearest neighbors approximation library *annoy* [21]. Both supervised and unsupervised versions of EmbedMNN algorithms were able to finish in just under 5 min, while Harmony took 5min30s plus an extra 30s of UMAP computation in order to obtain a 2D embedding. ScVI was the longest to complete with around 10 minutes in total, but in all fairness the minimum loss seemed to be reached between the 2 min and 3 min mark.

Computed joint representations were overall reasonable for all methods, with both convincing batch mixing and cell type clustering. Nonetheless, no method was able to provide both perfect batch mixing and meaningful cell type clustering, which is to be expected on such complex datasets (large number of cells, >10 patients, 10 cell types). Unsurprisingly, the supervised version of EmbedMNN outperformed all other methods by a large margin both in terms of local class purity and clustering class purity [Fig. 2c, 2f and Sup. Fig. 2], with a very low LISI-class score for all cell types and a near-100% cluster purity, as it leveraged complete label information. This allowed it to prune edges between cells of different types during the matching step, which results in a very clean cells graph to embed. On the other hand, supervised EmbedMNN is associated with poorer batch mixing [Fig. 2e], and clearer cluster delimitation which can for instance be an obstacle for some trajectory inference algorithms. The unsupervised version of EmbedMNN appears to be on par with the other methods, with reasonable LISI-class score and good LISI-batch score, as well as decent clustering purity [Fig. 2c,2e,2f and Sup. Fig. 2].

### Performing integration in gene space by using an appropriate embedding

In some applications, providing a joint embedding of datasets into an abstract space is not suited, as original features do carry important interpretable information. This is for instance the case when performing matrix factorization algorithms such as independent component analysis of non-negative matrix factorization. In this case, it is necessary to perform the integration directly within gene space, which brings some technical difficulties. Notably, gene spaces are typically very large which is detrimental for scalability of distance-based algorithms due to the curse of dimensionality. In this scenario, EmbedMNN, Harmony, scVI or BBKNN are not adapted, as they are unable to return an output in full gene space. We would normally need to find another integration method, install it, reprocess our data and adapt the workflow. In this example, we showcase how the modular nature of the *transmorph* model API can instead provide a way to easily adapt an existing model to a new application. To do this, we identify the embedding step of EmbedMNN not to be adapted to a full gene space application. To tackle this limitation, we can swap this module for something more adapted [Fig. 3b] like a linear correction step in gene space using correction vectors which is similar to what is used within MNN [4] and Seurat [8]. Given a reference dataset, the correction approach consists in first, finding some matchings between query and reference items, then compute correction vectors from these queries to their references, to finally propagate these correction vectors along the query dataset to end up with corrected profiles. This will demonstrate how simple module swapping allows reusing a model in a different scenario.

**Figure 3:**
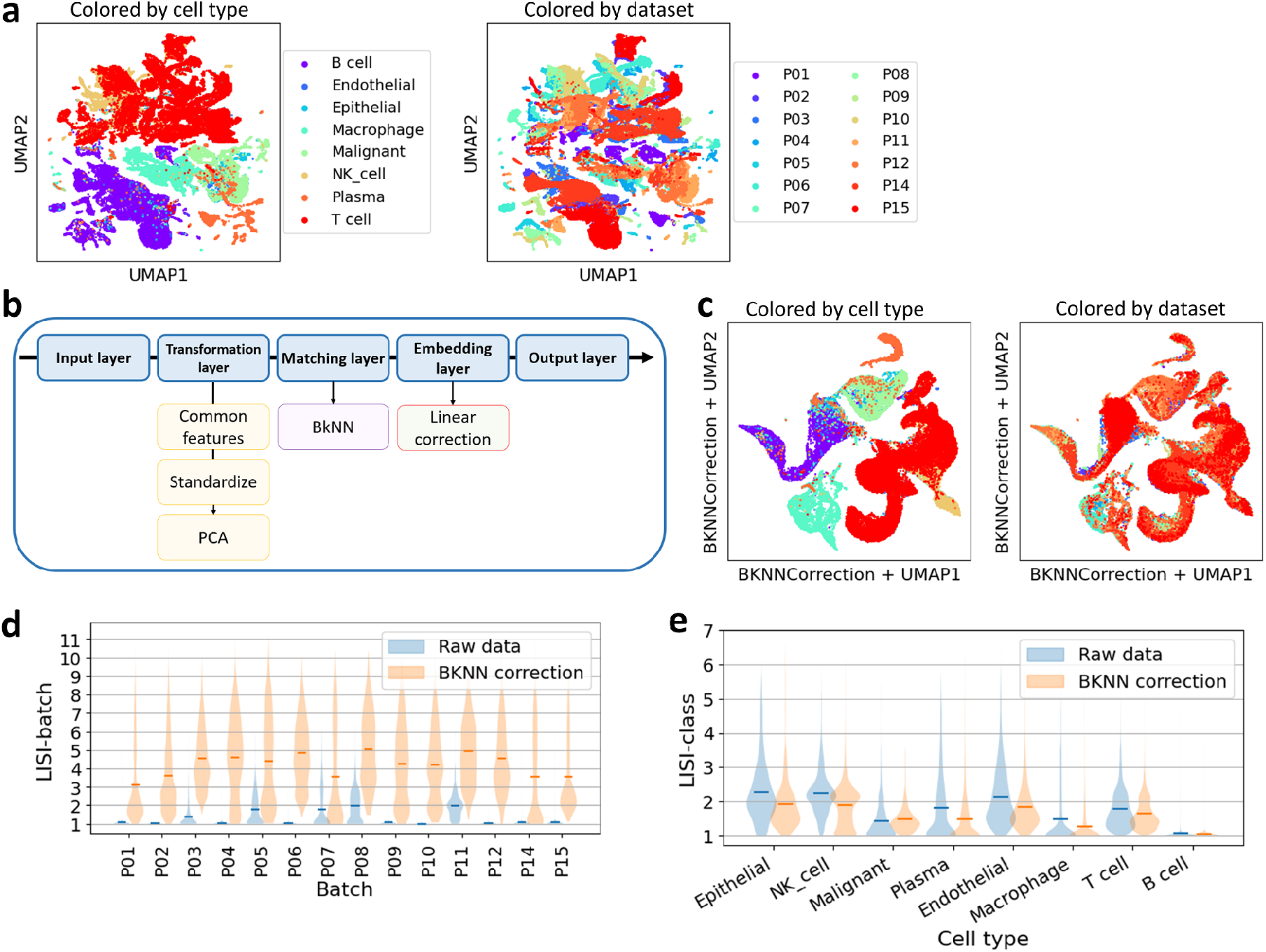
**a.** UMAP representation of the nasopharyngal carcinoma datasets (n > 60, 000) in their common genes space. **b.** Architecture of the *BKNNCorrection* integration model, computational modules are executed left to right and top to bottom. **c.** Integration result using *BKNNCorrection*. **d.** LISI-batch before and after integration (higher is better), mean is marked. **e.** LISI-class before and after integration (lower is better), mean is marked.

We use 14 nasopharyngal carcinoma datasets gathered from [22] to benchmark the strategy [Fig. 3a]. The goal is to embed these datasets in the space defined as the intersection of their common most variable genes, so that cells sharing the same annotation end up in close proximity after integration. This is once again a challenging task as the datasets are quite large (more than 60,000 cells to embed), there are 8 different cell annotations, some datasets do not contain cells from all types, and the embedding space is large for a geometrical approach (more than 900 genes). To measure integration quality from another angle, we carry out independent component analysis (ICA) on T cells from all datasets, which allows us to observe dataset-specific gene expression signals. As we can see, before integration the dataset-specific signal appears to be strongly correlated with several independent components (ICs) computed by ICA [Sup. Fig. 3, top].

BKNNCorrection completes in a very reasonable time of 1 minute and 33 seconds, and provides a convincing correction [Fig 3c, 3d, 3e]. We were not able to successfully carry out Seurat integration on these datasets in a reasonable time and memory usage. According to the LISI metric we use to measure integration quality, LISI-batch is greatly increased while LISI-class is maintained if not decreased after integration using BKNNCorrection. Overall, this showcases how *transmorph* provides a new way to easily tweak models, allowing them to tackle different scenarios with satisfying efficiency and integration quality. We also eventually ensure most of the dataset-specific signal has disappeared after integration [Sup. Fig. 3, bottom], resulting in a weak correlation with any of ICs recomputed by ICA on the integrated dataset. This is a desired property for subsequent accurate interpretation of the independent components through, for example, functional enrichment analysis.

### Carrying out label transfer to predict cell cycle phases

Cell cycle is one of the most fundamental biological processes through which biological cells grow and divide, but is yet to be fully understood. Single-cell transcriptomics offer great insight into its properties and dynamics, as gene expression regulation is a key factor for cell cycle progression. Gene expression modulation during cell cycle can be visualized and interpreted by looking at the so-called cell cycle plots. In these plots, each cell is reduced to a small set of coordinates (typically between 2 and 4 [23]), each of those corresponding to the average transcription activity of genes associated with a specific cell cycle signal (e.g. G1/S phase, G2/M phase, histones). In this configuration cells revolve along a one-dimensional looping trajectory throughout their progression in the cell cycle (see Fig. 4a). Studying the geometry of these trajectories and cell distribution along them can provide exquisite insight into cell cycle speed, cell growth or even eventual cell cycle arrest.

**Figure 4:**
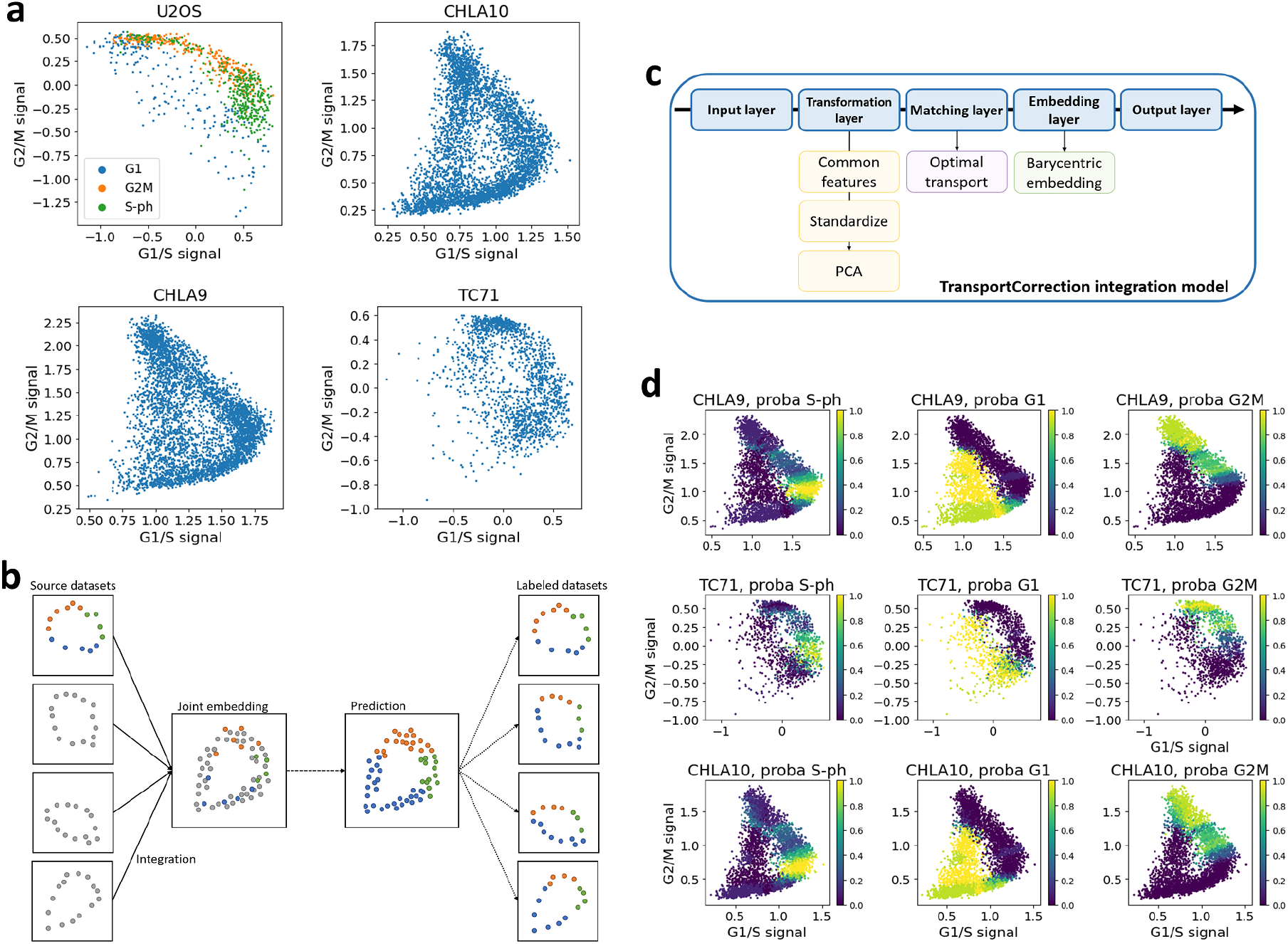
**a.** Osteosarcoma (U2OS) and Ewing sarcoma (CHLA9, CHLA10, TC71) datasets embedded in cell cycle space. U2OS cells are labeled with cell cycle phase. U2OS and TC71 shape in this space suggest they are examples of “fast” cell cycle. **b.** Label transfer strategy, with a first joint embedding step followed by a label prediction using a *k*-nearest neighbors classifier. **c.** Architecture of the TransportCorrection integration model. Computational modules are executed left to right and top to bottom. **d.** Label prediction probability for each query dataset (rows) and cell cycle phase (columns).

**Figure 5:**
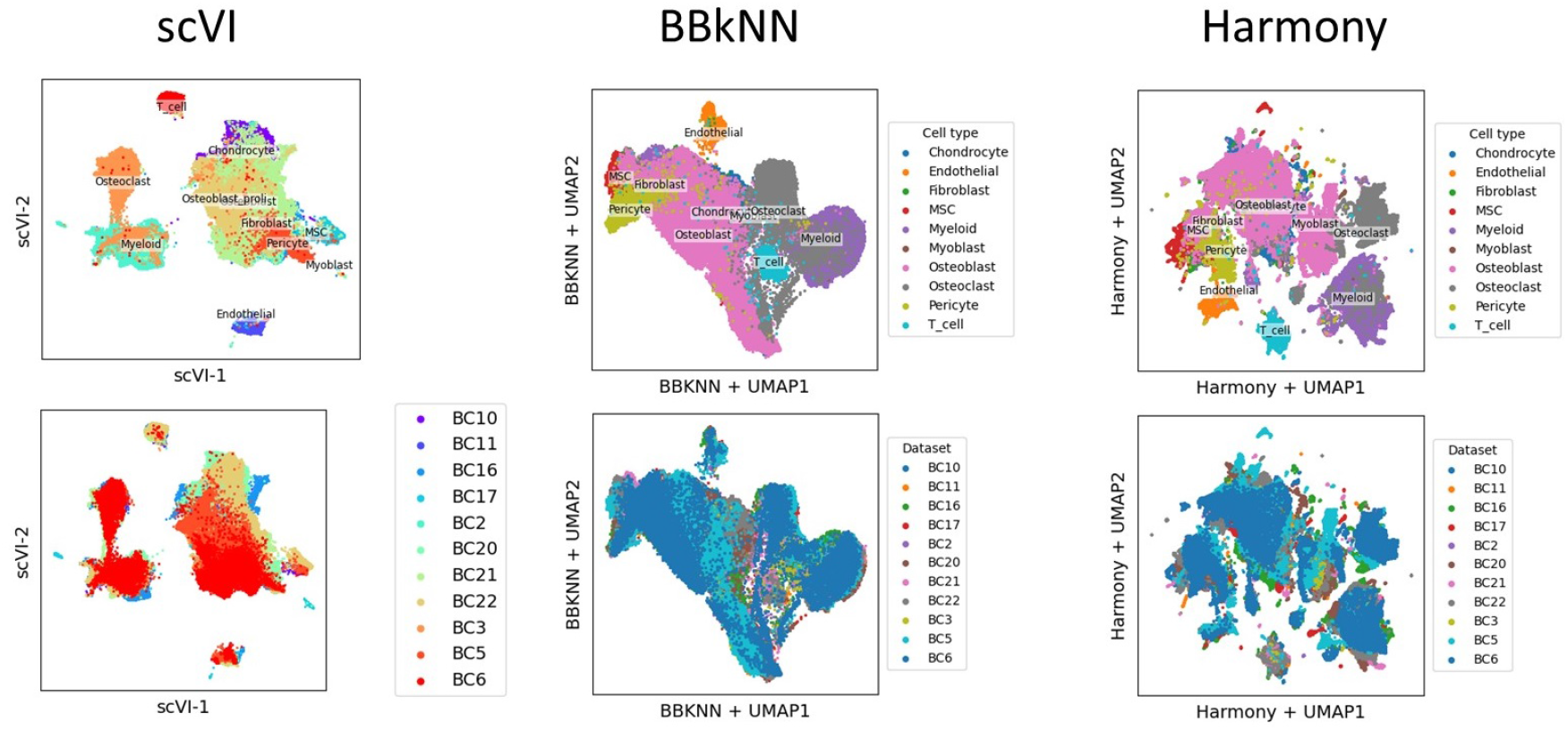
(Supplementary 1) Visualization of integration results between the different methods.

**Figure 6:**
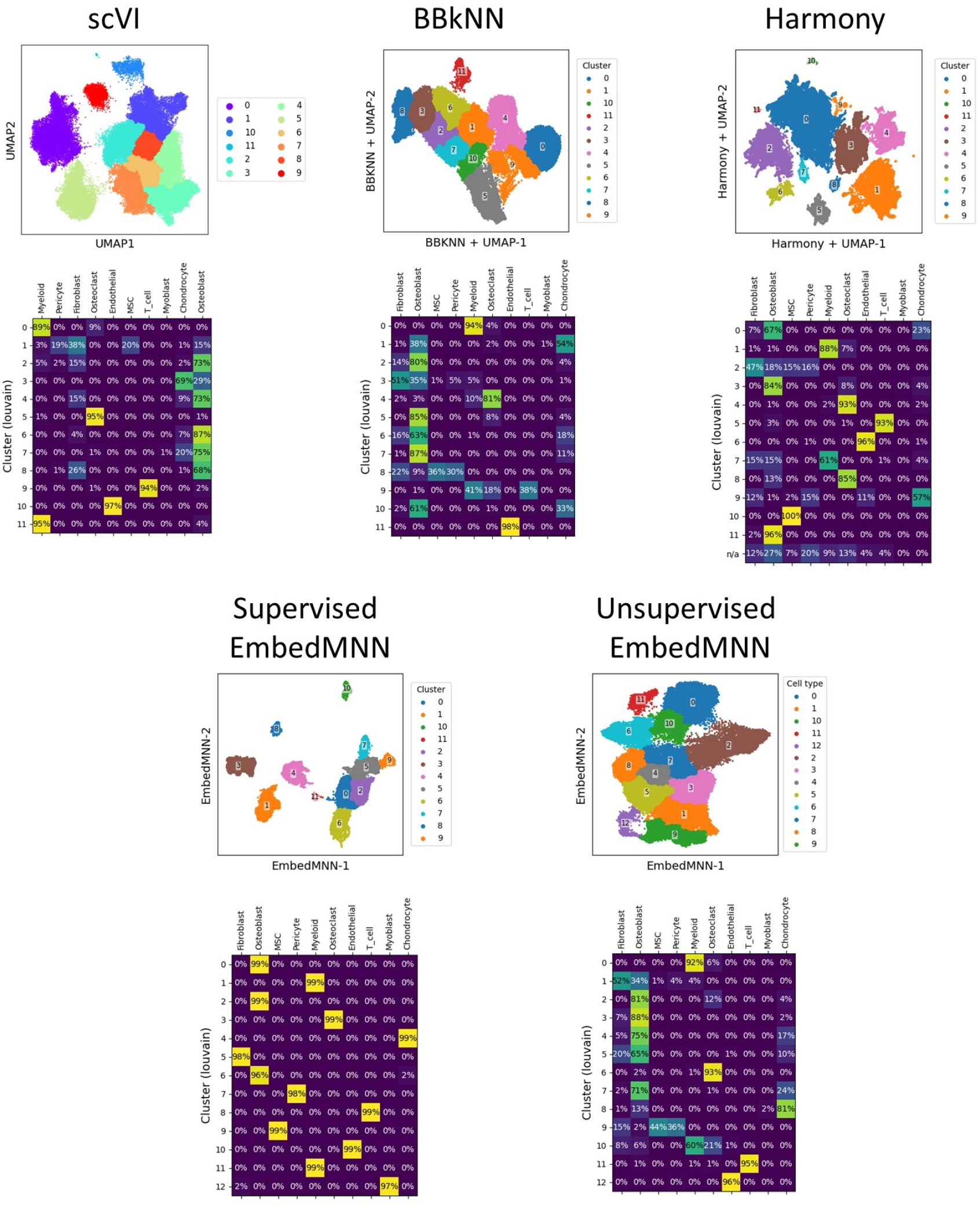
(Supplementary 2) Clustering purity results after integration. **Top:** Leiden cluster visualization. Resolution parameter was tuned to have an approximately equal number of clusters between methods. “Satellite” points in Harmony were ignored as they tended to generate a high number of very small clusters. **Bottom:** Cluster/cell type distribution, each value (*i,j*) corresponds to the proportion of cell type *j* in cluster *i*.

**Figure 7:**
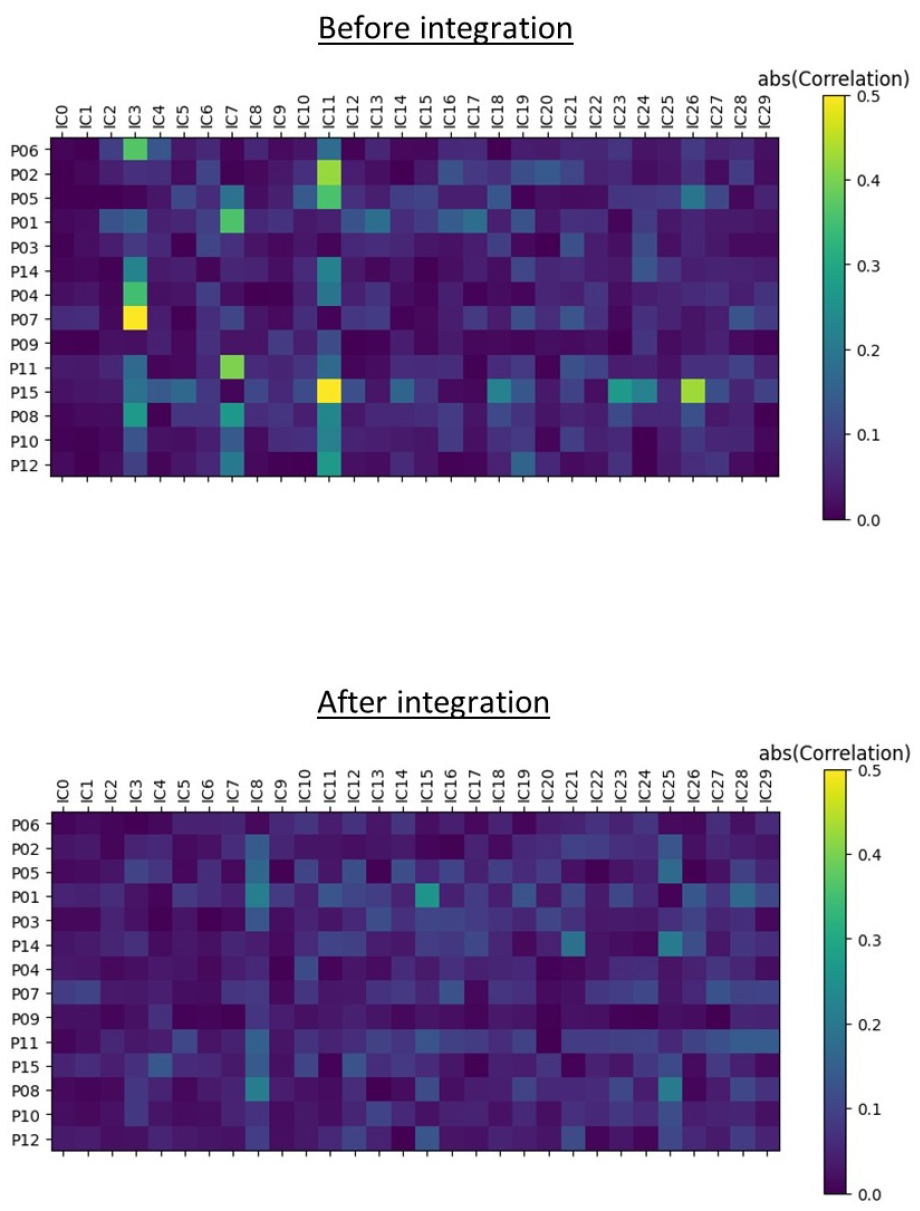
(Supplementary 3) Absolute correlation between independent component scores and original batches. We observe way weaker correlations between IC scores and batches after integration.

**Figure 8:**
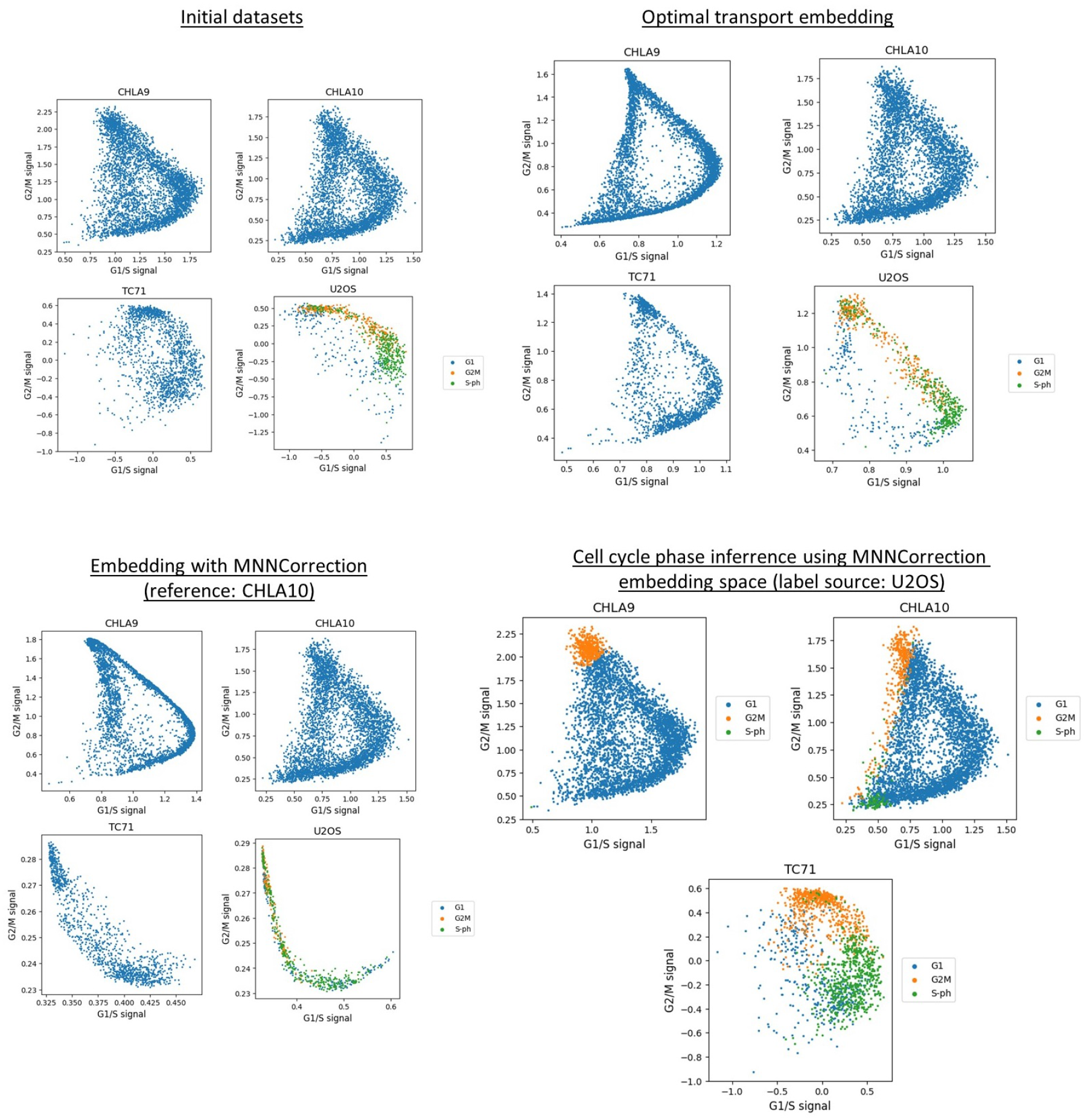
(Supplementary 4) Integration of cell cycle datasets for label transfer. *x* axis represents the average expression of G1/S phase genes, while *y* axis represents the average expression of G2/M genes.

A challenging question when studying cell cycle at the single-cell level is the automatic annotation of cells with cell cycle phases. It was studied experimentally in [24], where the authors used genetic constructs to follow the abundance of key cell cycle proteins which they can then relate to cell cycle phases, but doing so comes with important costs and experimenter time; a natural idea would be to transfer labels from datasets annotated using this methodology to other unlabeled ones. Unfortunately, this is not as easy as it seems: differences both in preprocessing, cell types and cell cycle properties can quite drastically affect a dataset topology and geometry, making many proximity-based methods irrelevant. A natural label transfer strategy can be pictured as follows [Fig. 4b]. First, we carry out data integration of all datasets into a common embedding space. Then, we predict cell cycle labels of unlabeled datasets in this common space using a supervised learning approach. Finally, the learned labels can be transferred back to the original representations to be interpreted.

In this experiment, we seek to automatically annotate three single-cell RNA-seq Ewing sarcoma datasets (CHLA9, CHLA10 and TC71) gathered from [25]. To do so, we will attempt to transfer cell cycle phase, author-provided annotations contained in an osteosarcoma dataset (U2OS) gathered from [24], onto the three Ewing sarcoma datasets. Preprocessing differences, geometrical specificities and apparent S/G2M label mixing within the U2OS reference dataset are tough difficulties to overcome both for integration and label transfer methods [Fig. 4a]. We first attempt to perform the integration using BKNNCorrection, setting CHLA10 as the reference dataset considering its good quality and representativity (cells are scattered uniformly around the trajectory, and the central “hole” is well resolved). Unfortunately, predicted cell cycle labels are not satisfying, almost all being predicted to belong to G1/S phases at any point of their trajectory [Sup. Fig. 4, bottom right]. Closer inspection reveals a very small number of matches across datasets using MNN, which can be caused by an absence of orthogonality between cell cycle factors and batch effects. This is a crucial hypothesis for neighbors-based dataset integration and its lack results in a poor matching quality, making integration unreliable.

Fortunately, there exist more appropriate algorithms for these situations: transportation-based matchings. They rely on discrete optimal transport theory that have been brought into the single-cell field a few years ago in [26], which can be pictured as looking for the most economical way to move mass in a metric space from a point cloud onto another. This class of problems yields a natural and harmonious way to match cells across batches, by operating at the dataset level instead of operating at the item level like MNN. We can use the *transmorph* pre-built model TransportCorrection inspired from SCOT [9] and Pamona [10], which consists in a few preprocessing steps followed by a transport-based matching, used to project every query item onto the barycenter of its matches [Fig 4c]. In this case we had to use the unbalanced formulation of optimal transport [27, 28] to account for cell cycle phase imbalance between ‘‘standard” and “fast” cell cycle datasets, which is implemented in *transmorph*. Label transfer using this model instead of BKNNCorrection yields a much better labeling, entirely interpretable and inline with the patterns we expect for “standard” and “fast” cell cycle [Fig. 4d]. This use case has demonstrated the need to choose an appropriate matching for a given scenario, which *transmorph* addresses.

## Discussion

Horizontal data integration and batch effect correction are key computational challenges, especially in computational biology to be able to properly analyze single-cell data from different batches or patients [1]. We expressed the need for modular methods to tackle this problem, and demonstrated the necessity to carefully combine trustworthy cell-cell similarity algorithms with relevant embedding algorithms. We also clearly showed how deceiving data integration can be when carried out improperly, which can be extremely detrimental for subsequent analyses. This alone motivates the need for more modular tools, where every algorithmic step can be controlled if necessary. To address this need and instead of introducing another data integration technique we present *transmorph*, a novel modular computational framework for data integration, implemented as an open-source python library. We provided a sound implementation for it, and demonstrated its value throughout various real-life applications both in terms of efficiency, quality and versatility.

We feel that increasing modularity and user agency often leads to bloated, over-engineered and impractical pieces of software. For this reason, we provide via *transmorph* several pre-built integration models ready to be used in daily workflows, with high efficiency and integration quality. For more advanced and specific applications, our framework also allows building integration models from scratch by combining a variety of algorithmic modules, all of which are implemented and optimized inside our library. We eventually provide complete interfaces which allow users to implement their own computational modules if they need to. All this is endowed with a rich software ecosystem including benchmarking datasets, integration metrics, monitoring and plotting tools.

We plan to continue maintaining *transmorph* in the future, in order to keep it up to speed with the ever-growing field of data integration methods. We will continue expanding it with new algorithms, either already existing or to come. We also would also like to add more support for vertical and diagonal integration, as for now the only diagonal matching is based on Gromov-Wasserstein which has a hard time scaling to the size of current data integration problems. For instance, we plan to use gene space transformation to deal with specific vertical integration cases such as integration between RNA-seq and ATAC-seq data. We would eventually like to add domain adaptation methods to our framework (for instance by including supervised PCA [29] or domain adaptation PCA [30] to our preprocessing steps), in order to tighten the bridge towards this growing research field which presents many similarities with data integration.

We are aware *transmorph* cannot be used in every data integration strategy: for instance, neural network-based or pure optimization approaches are not part of the framework yet, but these are things we will consider in the future. Also, there are still crucial questions to be answered in order to achieve high quality integration methods which we can really trust, especially in single-cell biology. Among these questions are the definition of relevant metrics to measure dissimilarity between cells (even more importantly across different domains), the research of sound and unbiased ways to measure integration quality and the necessity to continue to carry out exhaustive benchmarks to identify the most appropriate data integration methods and algorithms for a given use case. Last but not least, we believe *transmorph* is not the only possible way to bring modularity in the data integration fields, and we are looking forward to seeing new approaches tackling this ambitious challenge.

## Materials and methods

### Single-cell RNA-seq datasets

We use public datasets to benchmark our framework and compare its capabilities with other state-of-the-art integration pipelines. They were chosen to mimic various real-life scenarios, with total dataset sizes in the tens of thousands. All datasets contain RNA-seq data, acquired using 10X technology.

- The Zhou databank was collected from [18] through the Curated Cancer Cell Atlas (3CA) website, and contains osteosarcoma data from 5 different patients, ranging from 866 to 14,322 cells for a total of 64,557 cells. Each cell was annotated with a cell type among chondrocyte, endothelial, fibroblast, mesenchymal stem cell (MSC), myeloid, myoblast, osteoblast, osteoclast, pericyte, T cell.
- The Chen databank was collected from [31] using the 3CA website, and contains 61,870 nasopharyngeal cancer single-cell RNA-seq data from 14 different patients, ranging from 1,087 to 11,210 cells. Each cell was annotated with a cell type among B cell, endothelial, epithelial, macrophage, malignant, NK cell, plasma, T cell.

Raw counts have been preprocessed following standard guidelines using the *scanpy* python package [15]. First, cells with low gene counts or high mitochondrial gene expression were filtered. Raw counts were then normalized to 10,000 per cell, followed by neighborhood pooling using 5 nearest neighbors. Counts were then log(1 + *x*) transformed, and for each dataset the top 10,000 most variable genes were kept. All these preprocessed annotated databanks can be automatically downloaded through our framework, in order to serve for benchmarking integration methods.

### Cell cycle single-cell datasets and cell cycle signal

Raw counts for Ewing sarcoma cell lines datasets CHLA9, CHLA10 and TC71 were obtained from [25]. Raw counts and annotations for the osteosarcoma U2OS dataset were obtained from [24]. They were preprocessed according to state-of-the-art guidelines. Raw counts per cell were normalized to 10,000 to account for differences in global expression, and were then log(1 + x) transformed. Top 10,000 variable genes were kept in each dataset, and U2OS/TC71 values were scaled to unit variance/zero mean, clipped to −10/10 in order to simulate differences in preprocessing between datasets coming from different sources. Data points were eventually pooled, by setting every cell counts vector to the average of its 10 nearest neighbors (neighbors were determined using euclidean distance in a 30-PC space). We used cell cycle genes identified in [32] to characterize G1/S and G2/M signal. For fast cell cycle datasets TC71 and U2OS, we used only a subset of informative G1/S genes which helped to retrieve a proper loop signal (CDK1, UBE2C, TOP2A, TMPO, HJURP, RRM1, RAD51AP1, RRM2, CDC45, BLM, BRIP1, E2F8 and HIST2H2AC).

### Batch *k*-nearest neighbors matching

For *k* a positive integer, batch *k*-nearest neighbors is derived from the well-known *k*-nearest neighbors algorithm [33, 34]. It requires two batches **X***_a_* and **X***_b_* of respective size *n_a_* and *n_b_* to be embedded in a common metric space 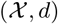, and works best when batch effect is orthogonal to biological signal of interest which seems to be a reasonable assumption in practice for most single-cell data, though it is challenging to verify. For every item **x***_a,i_* ∈ **X***_a_*, we denote by *r_a,i_*(*k*) the distance to its *k*-th nearest neighbor in **X***_b_*. We then define the batch *k*-nearest neighbors of **x***_a,i_* in **X***_b_* as 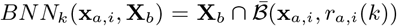 where 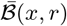 denotes the closed ball centered in 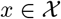 of radius 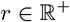. For every pair of samples **x***_a,i_* ∈ **X***_a_* and **x***_b,j_* ∈ **X***_b_*,

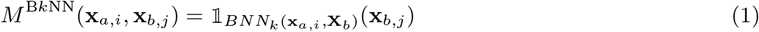

where 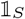 is the indicator function of set
*S*. Batch *k*-nearest neighbors derived algorithm have been successfully applied to dataset integration in the single-cell field, for instance in the BBKNN tool [5]. It tends to yield high quality matching results when the orthogonality of batch effect hypothesis is verified, and only needs the tuning of the *k* parameter (set at *k* = 15 in our applications). Furthermore, it also returns a much smaller number of edges compared to label matching, of the order of *kn_a_*, which greatly improves performance of subsequent pipeline steps. In terms of computational efficiency, its naive algorithm needs to compute the full *n_a_ × n_b_* distance matrix which corresponds to *n_a_n_b_* time and space complexity. This complexity being prohibitive in practice for large scale applications, we rather use nearest neighbors approximation schemes such as the nearest neighbor descent algorithm proposed in [35], which was found to run in quasi-linear time.

### Mutual nearest neighbors matching

For *k* a positive integer, *k*-mutual nearest neighbors [4] is an alternative to batch *k*-nearest neighbors. It tends to provide higher quality edges than batch *k*-nearest neighbors, at the cost of increased computation time and lesser edges number. The idea is to compute reciprocal batch *k*-nearest neighbors between two batches, and only keep the intersection of both edge sets. Given two batches **X***_a_* and **X***_b_* represented in a metric space 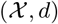 and two samples **x***_a_* ∈ **X***_a_* and **x***_b_* ∈ **X***_b_*, we first compute BNN*_k_*(**x***_a_*, **X***_b_*) and BNN*_k_*(**x***_b_*, **X***_a_*). Then,

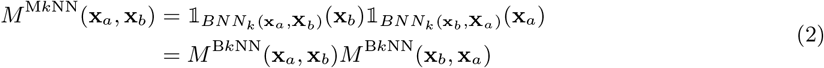

Mutual *k*-nearest neighbors tends to yield high quality matchings, inheriting all good properties from batch *k*-nearest neighbors, while also being symmetrical. It also works under the assumption that batch effect is orthogonal to biological effect, and always returns a smaller number of edges compared to batch *k*-nearest neighbors (*M*^*Mk*NN^(**x***_a_*, **x***_b_*) = 1 ⟹ *M*^*Bk*NN^(**x***_a_*, **x***_b_*) = 1). This fact often induces in practice the need to tune up the *k* parameter in order to have enough matching edges for subsequent pipeline steps to be stable.

### Graph embedding algorithm and clustering

Joint graph embedding is an algorithm able to build a joint weighted graph whose vertices are samples from all batches, and two samples are linked together if they appear to be similar. This graph being weighted, it can then be embedded in a low dimensional space using UMAP [16] or MDE [17]. The joint embedding algorithm consists of four major steps.

1. For each batch, compute its *k*-nn graph weighted according to UMAP membership methodology.
2. For each pair of batches, weight matching edges according to UMAP membership methodology.
3. Build a joint graph combining edges of steps 1 and 2, possibly selecting only the heaviest edges.
4. Embed the joint graph in an abstract feature space using a graph embedding optimizer such as UMAP or MDE.

Let 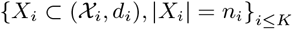 be a set of finite datasets to integrate, each expressed in a metric space. Let us also assume we are provided for every pair of datasets *X_i_* and *X_j_* a matching **M***_ij_* between *X_i_* samples and *X_j_* samples. We start by computing, for each batch *X_i_* its directed *k*-nearest neighbors graph *G_i_* = (*X_i_, E_i_*), weighted using UMAP methodology for computing membership strength [16] resulting in an adjacency matrix **K***_i_* describing a *k-nearest neighbors strength graph*. This guarantees every sample contains at least one edge of weight 1, and helps uniformize weights regardless of batch-specific point density; note that this graph is not symmetrized yet.

The next step is to convert all matching matrices to membership strength matrices so that edge weights are of the same nature as **K***_i_ k*-nearest neighbors graphs. For two batches *X_i_* and *X_j_* associated with a matching matrix 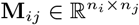, we distinguish two cases.

- If 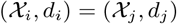 which can often be achieved between datasets of the same data type using common features selection, weights can be chosen using UMAP methodology on the bipartite matching graph **M***_ij_* using distance between matched points, yielding *matching strength graph* described by matrix **S***_ij_*.
- In the general case 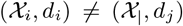, we can leverage matching strength contained in 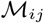 to produce a dissimilarity measure, for instance using the trick **D***_ij_* = inv(**M***_ij_* + 1) where inv(**M**) denotes coordinate-wise matrix inversion. This dissimilarity can then be used to compute the membership matrix as described in the previous case.

Once all *k*-nearest neighbors strength matrices **K***_i_* and matching strength matrices **S***_ij_* have been computed, they can be assembled in a joint graph *G* of all batches described by an adjacency matrix **G** whose blocks are defined as

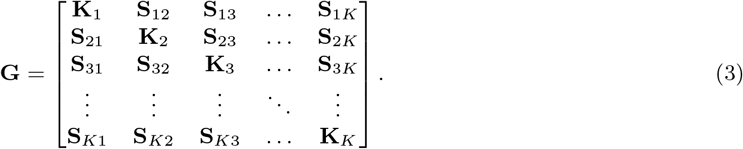

This matrix typically contains a very large number of edges, especially in large scale applications. This tends to increase convergence time of the graph embedding step. Furthermore, vertices tend to have a very variable number of edges in **G** which can result in embedding instability. To counterbalance these properties, we choose to first carry out an edge pruning step based on **G**. Given a target number of neighbors k_t_ > *K*, for every **G** row, all values below the *k_t_*-th largest are set to 0. **G** is eventually symmetrized into 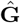 using

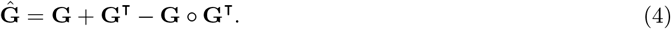

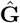 can eventually be embedded in an abstract feature space using a graph embedding optimizer such as UMAP [16] or MDE [17]. To perform clustering on this representation we carried out the Leiden algorithm [20] on this final representation, tuning the resolution parameter so that the number of significant clusters is between 10 and 15 in each case. on particular, Harmony had many “satellite” points which resulted in micro-clusters we decided to ignore in order to get an accurate comparison between methods.

### Local inverse Simpson’s index

Local inverse Simpson’s index (LISI) is an objective integration metric introduced in Harmony [3], which assesses neighborhood heterogeneity of a data point in terms of a given label. Simpson’s diversity index is a diversity metric notably used in ecology to measure class diversity in a set of objects by computing the probability for two randomly selected items to share the same class. We have chosen to simplify LISI in our implementation by removing the custom UMAP-like cell-cell weighting introduced in Harmony in order to increase its computational performance, and make it less dependent on local geometry. For any set of objects *S* = {*x_i_*}*_i≤n_* endowed with labels 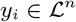, we denote by *n_l_* for 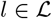 the number of samples in *S* with label *l*. Then, Simpson’s index of set S is given by

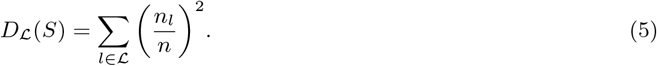

For *k* > 0 a *perplexity* parameter (we use *k* = 90) and x an embedded point, we compute its *k*-nearest neighbors *k*-nn(*x*) which is used as set *S*. 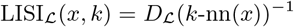 is defined as the inverse of Simpson’s index and estimates, for a given embedded point x, label diversity in its *k*-nearest neighborhood. As suggested in Harmony, we can use LISI in two modes:

- Batch-LISI (bLISI), where points are labeled by their initial batch. This metric measures local batch diversity in embedding, higher diversity being better.
- Class-LISI(cLISI), where points are labeled by their class. This metric measures local class diversity in the embedding, lower diversity being better.

Monitoring these two values allows objectively comparison of different integration pipelines, ideally increasing bLISI and decreasing cLISI.

### Linear correction

Inspired from Seurat [8], linear correction is a linear merging based on first computing a set of correction vectors, and then use them to correct batches with respect to a reference batch **X***_r_*. It not only (1) necessitates to choose a reference batch, but also (2) for all batches to be embedded in a common feature space. It can work with *incomplete* matchings, meaning not every cell in the query batch necessitates to be matched with a cell in the reference batch. To correct a given batch, the algorithm follows a two-step process: it first computes correction vectors from matched samples to reference samples, and then extrapolates the correction vectors to the unmatched samples.

Let **X**_1_,…, **X***_K_* be *K* batches each represented in a common features space 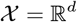, respectively containing *n*_1_,…, *n_K_* samples. Let 1 ≤ *r* ≤ *K* be the reference dataset index, and 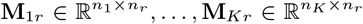 be *k* matching matrices between each batch and the reference - by convention **M***_rr_* = **I***_n_r__*. Each batch **X***_s_* is corrected independently towards the reference **X***_r_*. Let 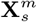 be the matched samples of **X***_s_* (**X***_s_* rows so that corresponding row in **M***_sr_* contains at least one nonzero element), and 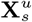 be the unmatched samples (the other rows). The first step is to compute the projection of each matched sample to its barycenter 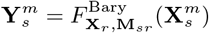. For every matched sample, we compute each correction vector

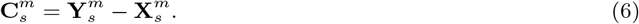

Total correction vectors **C***_s_* are then computed as follows. For every matched cell, we set the correction vector to the corresponding 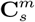 entry. Otherwise, we set the correction vector to the one of its closest matched cell along edges of a *k*-nearest neighbors graph. If there is no matched cell within the cell’s connected component, we consider this may be a specific cell type with no corresponding cell in the reference dataset, and it is left uncorrected. There exist variants for this step, for instance averaging correction vectors among sets of points (e.g. neighborhood or clustering) instead of selecting just one in order to smooth the final representation. In the end, merging is simply performed as

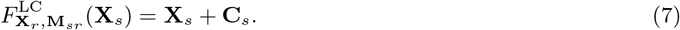

### Independent component analysis

Independent component analysis (ICA) is a matrix factorization (MF) approach where the signals captured by each individual matrix factors are optimized to become as mutually independent as possible. ICA was shown to be a useful tool for unraveling the complexity of cancer biology from the analysis of different types of omics data. Such works highlight the use of ICA in dimensionality reduction, deconvolution, data pre-processing, meta-analysis, and others applied to different data types (transcriptome, methylome, proteome, single-cell data) [36].

In ICA we search for an approximation of the observed probability density function *P*(*x*_1_, *x*_2_,… *x_n_*) by 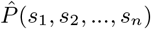, where new *s_i_* variables are some linear combinations of the initial variables *x_i_*. We search for such linear transformation that 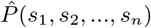 deviates as little as possible from the product of its marginal distributions *P*(*s*_1_) × *P*(*s*_2_),… *P*(*s_n_*) where the deviation is usually defined in terms of information geometry (e.g., as Kullback-Leibler divergence). It is shown that ICA is efficient in detecting and correcting the batch effects in omics datasets [36].

ICA is not a data dimensionality reduction technique *per se*: therefore, it is usually applied on top of reduced (e.g., by standard PCA) and whitened representation of the initial dataset. Therefore, the choice of the number of independent components is an important hyperparameter [37]. In the simplest approach, ICA solution represents a rotation of the whitened data point cloud such that each normalized coordinate deviates as much as possible from the standard Gaussian distribution [38].

In our experiments we used the stabilized version of ICA [39] which is shown to be the optimal MF approach for reproducible analysis of transcriptomic data [40].

### Discrete optimal transport

Discrete optimal transport (OT) problem can be naturally pictured as follows [28]. Assuming a set of n warehouses containing goods to deliver to *m* factories, the optimal transport problem consists in finding the cheapest way to transport all goods to factories knowing cost of transporting goods is proportional to both mass carried and distance traveled. Originally brought into the field as a way to predict cell fate [26], it has more recently been shown to be an interesting asset for matching cells across datasets in integration tools like SCOT [9] and Pamona [10].

Formally, let 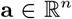 and 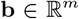 be two histograms, meaning **a** and **b** coefficients are non-negative and each vector sums up to 1. In our analogy, **a** represents the amount of goods stored in each warehouse, and **b** the capacity of each factory. We are provided a cost matrix 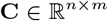, where **C***_ij_* is the cost of moving one unit of mass from *a_i_* to *b_j_*. The optimal transport problem from **a** to **b** given cost **C** can then be expressed as follows:

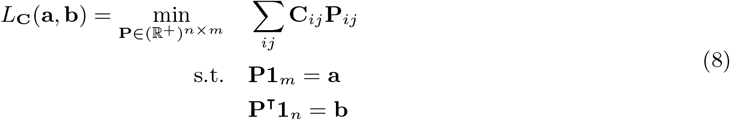

*L*_C_ (**a**, **b**) is called the *Wasserstein* distance between **a** and **b** for transport cost **C**, and is the cheapest cost to transport all mass from **a** to **b** in this setup. The optimal *transport plan* **P**\* is the valid **P** minimizing **Eq. 8,** and can be row-normalized to **1**_n_ to be used as a probabilistic matching between **a** and **b**.

In practice, optimal transport can be computed between two datasets 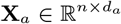 and 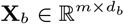 with vectorized samples in row. In this case, it is common to define 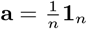 and 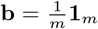, and to transform **X***_a_* and **X***_b_* so that they are expressed in the same feature space *d*. It can be achieved for instance by selecting the genes intersection, yielding 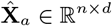 and 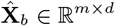. The cost matrix 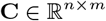 is then typically defined as the pairwise distance matrix between 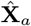 and 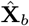, for a given distance. If **X***_a_* and **X***_b_* cannot easily be embedded in a common features space, the Gromov-Wasserstein approach is in general a better alternative. We use here optimal transport as a matching algorithm, by considering the row-normalized 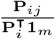 as the probability that cell *i* from dataset **X***_a_* is similar to cell *j* from dataset **X***_b_*.

There are obvious limitations to this procedure, notably the mass conservation issue: OT will always move *all* mass from **a** to **b**, regardless of the possible batch-specific samples. Consequently, all **X***_a_* cells will be mapped to at least one cell in **X***_b_*, even though some of these **X***_a_* cells may be of a cell type missing in **X***_b_*. Even worse, if there is a class imbalance between dataset (e.g. 50% of cell type A in dataset **X***_a_*, and 25% of cell type A in dataset **X***_b_*), there will necessarily be wrong assignments using this method. Exact computation of optimal transport is furthermore computationally expensive, of the order of 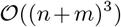 which makes it inefficient for large scale problems (typically above 10^4^ points). The next section presents an alternate formulation which provides a good approximation of the solution at a more reasonable cost, and subsampling schemes can also be used.

### Entropy-regularized unbalanced optimal transport

The optimal transport problem can be approximated using an additional entropy term [41, 28], which allows the minimization to be carried out using an efficient iterative procedure. For a given transport plan **P**, its entropy is defined as

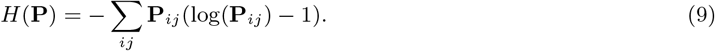

*H*(**P**) is 1-strongly concave given its Hessian *∂*^2^*H*(**P**) = –diag(1/**P***_ij_*) and **P***_ij_* ≤ 1. –*H*(**P**) can then be used as a regularizer term in Eq. 8 with a regularization term *ε* > 0 [42], making the objective *ε*-strongly convex:

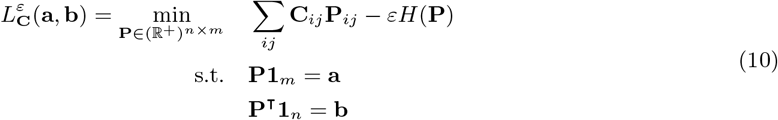

Sinkhorn-Knopp algorithm can be used to optimize the objective, we invite the reader to refer to [41] for details. In short, the goal is to decompose a transport plan **P** = diag(**u**)**K** diag(**v**), where **u** and **v** are the unknown *scaling variables* and **K** can be derived from the parameters. **u** and **v** can be approached using an iterative two-step normalization procedure. The smaller ε, the closer the objective is from unregularized formulation, at a cost of decreased convergence rate. According to [43] and assuming n = *m* for simplicity, this algorithm computes a τ-approximate solution of the original optimal transport problem in 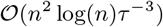 operations. It allows tackling larger scale problems in reasonable time. In practice, the resulting transport plan is often more fuzzy and less sparse than the exact solution, which necessitates filtering small values to stay efficient.

As stated previously, one of the major drawbacks of optimal transport is its constraint to always move all mass from source distribution to target distribution. As there is almost always class imbalance between single-cell datasets, this hard constraint necessarily causes matchings between cells of different cell types. This bad property can be worked around using an alternative unbalanced optimal transport problem [27]. The idea is to relax the hard mass conservation constraint, by rather penalizing mass discrepancy via a divergence *D_φ_*. Given two penalty coefficients *τ*_1_ and *τ*_2_, the objective function is written as

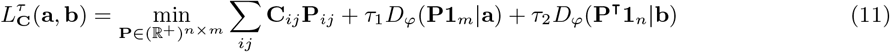

This objective function can also be optimized using an adaptation of the Sinkhorn-Knopp algorithm. Removing the hard mass conservation constraint helps in practice to deal with situations of class imbalance between datasets, while staying entirely unsupervised.

### Barycentric embedding and label transfer

Barycentric merging is the simplest merging to set up. It works under three assumptions, (1) one batch **X***_r_* is defined as *reference* and all batches will be corrected towards it; (2) reference batch **X***_r_* must be expressed in a feature space; (3) for every matching 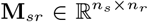, every row must have at least one nonzero element (∥**M***_s,r_***1***_nr_*∥_0_ = *n_s_*). Assumption (1) is fulfilled by user choice, and necessitates choosing a good quality batch with representative items within every sample type. Reference choice always introduces a bias in the integration, which should not be overlooked in results interpretation. Assumption (2) is easy to verify in practice, as datasets are often vectorized and represented as *n* × d real valued matrices. Assumption (3) necessitates to choose a *semicomplete* matching, which maps every sample from batch *X_i_* to at least one sample from batch **X***_r_*. Transportation-based matchings usually verify this assumption, while nearest neighbors-based matchings usually do not. Failing to verify assumption (3) will cause non-matched points to be projected to the 0 of **X***_r_* feature space.

Let *X_s_* be a batch to correct with respect to the reference batch **X***_r_* given a semicomplete, row-normalized matching matrix **M***_sr_*. For every sample *x_k_* ∈ *X_s_*, the *k*-th row *α_k_* = **M***_sr,k_*, provides a weighting vector which assesses the likelihood of *x_k_* corresponding to any sample of **X***_r_*. Barycentric merging *F*^Bary^ will then project *x_k_* into **X***_r_* feature space 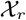 so that

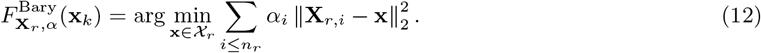

It is easy to show **x***_k_* = ∑*_i≥n_r__* α*_i_***X***_r,i_*, is the solution to this problem. Therefore, *F*^Bary^ can be easily generalized to project the whole X_s_ dataset onto **X***_r_* given **M***_sr_* via

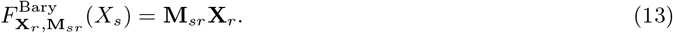

Barycentric merging has been used in several data integration pipelines such as Seurat [8], SCOT [9] and Pamona [10], and generally yields good results, though there are a few downside to consider. First, choosing a reference introduces a high bias in the integration, and in some applications there may be no good option for reference; for instance, every batch could miss at least one sample class. The barycenter problem also intrinsically relies on a metric. This is an issue for high dimensional problems, for instance in scRNA-seq datasets where curse of dimensionality is a real concern; in this case, barycenter has little to no interpretable sense. A common solution is to first reduce dimensionality of **X***_r_*, using component analysis or non-linear methods such as UMAP [16] or MDE [17]. One of the other uses of this method is to use a matching computed in a different space than the final embedding. Typically, one computes a matching in a lower dimensional representation (e.g. PC space), but uses total feature space for the embedding. This notably allows obtaining corrected feature counts for all batches with respect to a reference. Combined with a high quality matching and reference batch, barycenter merging can nonetheless provide an efficient, high quality integration without necessitating batches to be originally in the same space.

Label transfer was carried out in the integrated space using a simple nearest-neighbors approach. We used the `scikit-learn` implementation of a *k*-nearest neighbors classifier using *k* =10 and euclidean distance. We decided to assess label probability for a given cell x and label *l* by the frequency of *l* in x’s neighborhood.

### Benchmarking methods and parameters

All benchmarks have been run on a laptop equipped with 32GB of RAM, an Intel CPU i7-10750H (12 cores) processor at 5GHz and a NVIDIA GPU GeForce GTX 1650 Ti Mobile.

- EmbedMNN was used on preprocessed counts with *transmorph* v0.2.0, using default parameters: “bknn” matching, 10 matching neighbors, 10 embedding neighbors, UMAP optimizer and 2 dimensions.
- BKNNCorrection was used on preprocessed counts with *transmorph* v0.2.0, using default parameters: “bknn” matching, 30 matching neighbors, 10 linear correction neighbors.
- TransportCorrection was used on preprocessed counts with *transmorph* v0.2.0, using solver=“unbalanced”, entropy _epsilon=0.02, unbalanced_reg=5.
- Harmony was used with default parameters directly on preprocessed counts using the rpy2 python interface. We also tried the `harmonypy` python implementation, interfaced *via* `scanpy`. It successfully converged in under 10 iterations in both cases, and produced comparable results.
- scVI was used on preprocessed counts following authors guidelines with n_layers=2 and n_latent=30, and was optimized during 124 epochs (automatically chosen by the software).
- We used the `scanpy` implementation of BBKNN on preprocessed counts. We carried out BBKNN with default parameters on a 50-PC representation of datasets using default parameters, using neighbors_within_batch=3 and 10 annoy trees.
- Seurat was used in RStudio after converting AnnData datasets to h5seurat using the SeuratDisk package. We carried out the integration using SelectIntegrationFeatures, FindIntegrationAnchors and IntegrateData with default parameters. We were not able to complete the last integration step despite our efforts due to memory usage issues.

## Code and data availability

*transmorph* framework is available at https://github.com/Risitop/transmorph, and can also be downloaded from the PyPi repository (version 0.2.4 at time of writing). Datasets can be directly downloaded from the package, and scripts to generate figures can be found in supplementary materials.

## Author contributions

A.F., L.C and A.Z conceptualized the package design and the computational study. A.F. and L.C. developed the package, A.F. carried out the applications, A.F, L.C., O.D. and A.Z wrote the manuscript, A.Z. and O.D. directed the project.

## Acknowledgements

We thank Alexander Chervov (Institut Curie, U900) for his suggestions on how to process TC71 and U2OS fast cell cycle datasets, Marianyela Petrizzelli (Institut Curie, U900) for her help in testing *transmorph* on non-Linux systems, Jane Merlevede (Institut Curie, U900) for testing *transmorph* at an early stage, Vicent Noël (Institut Curie, U900) for his suggestions on how to properly package and test *transmorph* and in alphabetical order Jonathan Bac (Institut Curie, U900), Nicolas Captier (Institut Curie, U900), Marco Ruscone (Institut Curie, U900) and Julien Vibert (former Institut Curie, U830) for the many insightful discussions and suggestions around this project.

## Funding

This project has been financially supported by French government under management of Agence Nationale de la Recherche as part of the “Investissements d’avenir” program, reference ANR-19-P3IA-0001 (PRAIRIE 3IA Institute) and by European Union’s Horizon 2020 program (grant No. 826121, iPC project).

## Conflict of interest

A.Z. is currently an employee of EvoTec.

## Notes

### Competing Interest Statement

Andrei Zinovyev is an employee of the Evotec company.

